# Gait kinematics dynamically tune optic flow motion-in-depth perception during human locomotion

**DOI:** 10.64898/2026.09.24.754134

**Authors:** Edward A B Horrocks, Jocelyn Y L To, Isabelle Mareschal, Aman B. Saleem

## Abstract

Behavior generates continuous sensorimotor contingencies that the brain must process. In particular, locomotion produces optic flow, which is frequently noisy and ambiguous during navigation. Integrating non-visual self-motion signals can therefore provide a computational advantage to resolve this sensory uncertainty. To investigate this we developed a novel head-mounted 3D motion coherence paradigm. We found that walking dissociably alters perceptual sensitivity and bias for optic flow, dependent on stimulus speed. Rather than acting as a binary state change, these modulations scale dynamically with step frequency. While slow stepping impairs sensitivity to optic flow, fast stepping rescues this deficit. Simultaneously, higher step frequencies drive a progressive shift in perceptual bias, consistent with canceling expected self-generated flow. Our results indicate that optic flow perception during locomotion is modulated by active processes, encompassing both discrete behavioral state changes and continuous, dynamic sensorimotor integration. Broadly, these findings demonstrate that the brain leverages natural contingencies between behavior and expected sensory feedback to actively modulate perception.

**Highlights:**

- Head-mounted 3D motion coherence task probes optic flow perception while walking.
- Walking dissociably alters optic flow sensitivity and bias, dependent on stimulus speed.
- Optic flow sensitivity and bias covary with observer step frequency.
- Slow stepping impairs sensitivity, but fast stepping restores or enhances it.

## Introduction

An animal’s behavior generates continuous sensory feedback, resulting in a set of tightly coupled sensorimotor contingencies. Deciphering how the brain processes these contingencies is fundamental to understanding how animals interact with their environment. A striking example of this is optic flow, the full-field visual motion that results from locomotion ^1–3^. The co-evolution of locomotion and visual motion perception indicates that the joint neural processing of these variables is a highly ubiquitous feature observed across the animal kingdom^1,4–8^. In idealised environments, optic flow provides unambiguous information about an animal’s self-motion^9^. However, in real-world environments different three-dimensional structural features generate complex, non-uniform optic-flow patterns, which alongside sensory noise such as independently moving objects, render optic flow inherently ambiguous. Consequently, there are computational advantages to integrating non-visual cues for self-motion with optic flow signals to resolve sensory uncertainty ^10–13^.

Historically, a prominent viewpoint posited that the brain attenuates self-induced visual motion signals to ensure perceptual stability of the environment ^13–15^. However, growing evidence suggests that locomotion engages a more active, adaptive process that optimises the encoding of behaviorally relevant visual inputs ^16–18^. For example, across multiple mammalian species, locomotion is associated with the preferential encoding of the visual periphery ^19–21^, where optic flow vectors are generally largest and most informative about self-motion. Moreover, neurons that encode optic flow also often encode non-visual self-motion signals ^22–24^, indicative of integrative neural processing. Accordingly, visual and non-visual cues may undergo mandatory perceptual fusion in humans, preventing conscious access to optic flow in isolation ^25,26^.

Consistent with active sensorimotor processing, psychophysical studies demonstrate that human optic flow perception changes during locomotion ^16,17,27–30^. To date, these sensorimotor interactions have primarily been characterized through the lens of visual speed estimation or discrimination, in idealized, noise-free environments. While these studies have revealed that perceptual bias and sensitivity for optic flow can be altered during locomotion, real-world navigation presents a more demanding computational challenge: the brain must resolve the three-dimensional direction of motion-in-depth while continuously integrating noisy, ambiguous sensory evidence from across the visual field ^31–36^. Determining how this integration of evidence is modulated by locomotor state and the continuous kinematics of natural gait is important for understanding the extent to which navigation through noisy environments relies on isolated sensory processing or more integrated sensorimotor mechanisms.

To understand the nature of this sensorimotor coupling, we developed a novel 3D motion coherence task to investigate how human locomotion and its underlying gait kinematics alter the perception of optic flow motion-in-depth. Utilizing a wireless head-mounted display with an open-loop design, we isolated internal non-visual signals by keeping visual inputs strictly independent of observer movement. We found that locomotion dissociably alters participants’ perceptual bias and sensitivity dependent on optic flow speed, arguing against generic state-change effects like increased cognitive load. Crucially, by leveraging natural intra-participant variance in step frequency, we demonstrate that perceptual sensitivity and bias are not fixed traits of the walking state, but dynamically covary with ongoing gait kinematics. Taken together, our findings demonstrate that the brain leverages the natural rhythm of an animal’s behavior to actively calibrate visual perception.

## Results

### A 3D motion coherence paradigm to investigate optic flow motion-in-depth perception during different locomotion states

To investigate how non-visual signals of locomotion influence optic flow perception, we developed a novel 3D motion-in-depth direction discrimination task based on classical motion coherence paradigms. Participants performed a two-alternative forced-choice (2-AFC) task to identify the global motion direction of a cloud of moving spheres moving either ‘towards’ or ‘away’ from the observer viewpoint (Supplementary Movie 1). Participants performed the task in trials while standing still or walking (‘Stationary’ and ‘Walking’ states; Figure 1a; Supplementary Figure 1). Stimuli were presented stereoscopically via a wireless head-mounted display, utilising an open-loop rendering design to ensure that visual stimulation remained consistent across both locomotion states.

**Figure 1:**
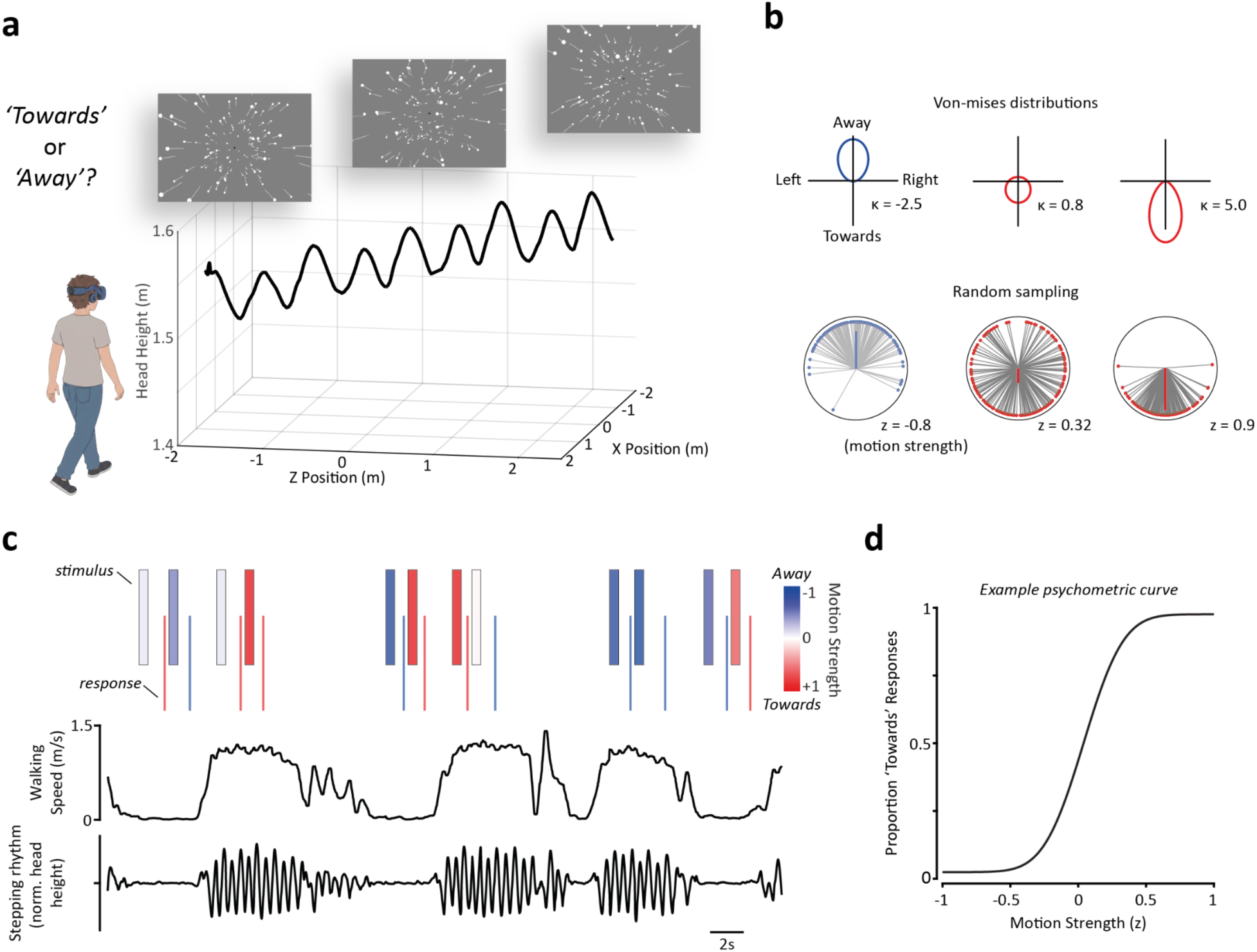
A 3D motion coherence paradigm to investigate optic flow motion-in-depth perception during different locomotion states. a) Overview of the experimental setup. Participants performed an optic flow motion-in-depth direction discrimination task while standing still and walking over solid ground. Optic flow visual stimuli (sphere fields) were presented via a wireless head-mounted display. Each trial block consisted of two consecutive stationary and two consecutive locomotion trials, randomly ordered. b) Example stimulus distributions for three trials. *Top Row*: Von Mises distributions shown in polar plot form. Blue polar plots indicate an ‘Away’ trial (μ=0) and red plots indicate ‘Towards’ trials (μ = π). κ values are the concentration parameter which determines the circular variance of each distribution. *Bottom Row*: Vector plots corresponding to the distributions above. Thin grey lines with coloured circles represent individual motion directions sampled from the corresponding Von Mises distribution above. The thick coloured lines represent the transformed stimulus value (z) used to perform data analysis - sampled stimulus motion vectors were projected onto the depth axis and then mean-averaged to produce the average motion-in-depth vector of the stimulus (values range from −1 to 1). c) Stimulus and behavioural data timeline from example trial blocks. *Top row*: Stimulus periods (boxes) and response times (lines; times represent the moment the verbal response from the participant was entered by an experimenter). Boxes and responses are coloured according to the motion strength of the trial and response given, respectively. *Middle and bottom rows:* Walking speed (middle row) and detrended head height of the participant. The rhythmic oscillation of head height was used to extract step frequency. d) An example psychometric curve showing the probability of a participant giving ‘towards’ response as a function of motion strength (z).

The stimulus consisted of a cloud of 3D moving spheres, an extension of random dot kinetogram (RDK) designs. At the onset of each trial, 175 white spheres appeared at random 3D coordinates and moved with a constant velocity along the azimuthal plane, whilst maintaining a fixed elevation, thus moving parallel to the floor in random directions. The motion directions of individual spheres were sampled from a Von Mises distribution with a mean (*μ*) facing either directly ‘towards’ or ‘away’ from the participant (Figure 1b). By manipulating the concentration parameter (*κ*) of this distribution, we systematically varied the global motion strength (analogous to motion coherence), and thus level of sensory uncertainty, with larger concentration parameters resulting in narrower distributions of motion directions and higher motion strength (Figure 1b). To determine if the perceptual effects of locomotion depend on the speed of optic flow, the stimulus was presented at one of three speeds - slow (15 virtual units (vu)/s), medium (50 vu/s), or fast (100 vu/s) - which varied randomly on a trial-by-trial basis. Locomotion conditions were interleaved in blocks of 4 trials, with each block consisting of a pair of stationary trials followed by a pair of walking trials, or vice versa (Figure 1c). As expected, participants’ perceptual judgements scaled reliably with motion strength, yielding characteristic sigmoidal psychometric curves (Figure 1d).

### Locomotion exerts dissociable, optic flow speed-dependent effects on perceptual sensitivity and bias

To evaluate how locomotion state and stimulus speed modulate optic flow motion-in-depth perception we used a Generalized Linear Mixed-Effects (GLME) model. We fit the model to single-trial binary responses pooled across participants (N = 16 participants; n = 12,068 total trials). We modeled the probability of a ‘towards’ response as a function of signed motion strength (Figure 1b,d), behavioral state (stationary or walking), and stimulus speed (slow, medium or fast) using a probit link function (Figure 2a,b). This approach allowed us to simultaneously characterize shifts in perceptual bias (captured by the model intercept, *β*_intercept_) and sensitivity (captured by the model slope, *β*_slope_ i.e., the motion strength weight; Figure 2c). By including interaction terms between behavioral state and stimulus speed, we were able to determine whether any locomotion-induced changes in perception were optic flow speed-dependent. To account for inter-participant variability in baseline task performance, we included random intercepts and slopes grouped by participant.

**Figure 2:**
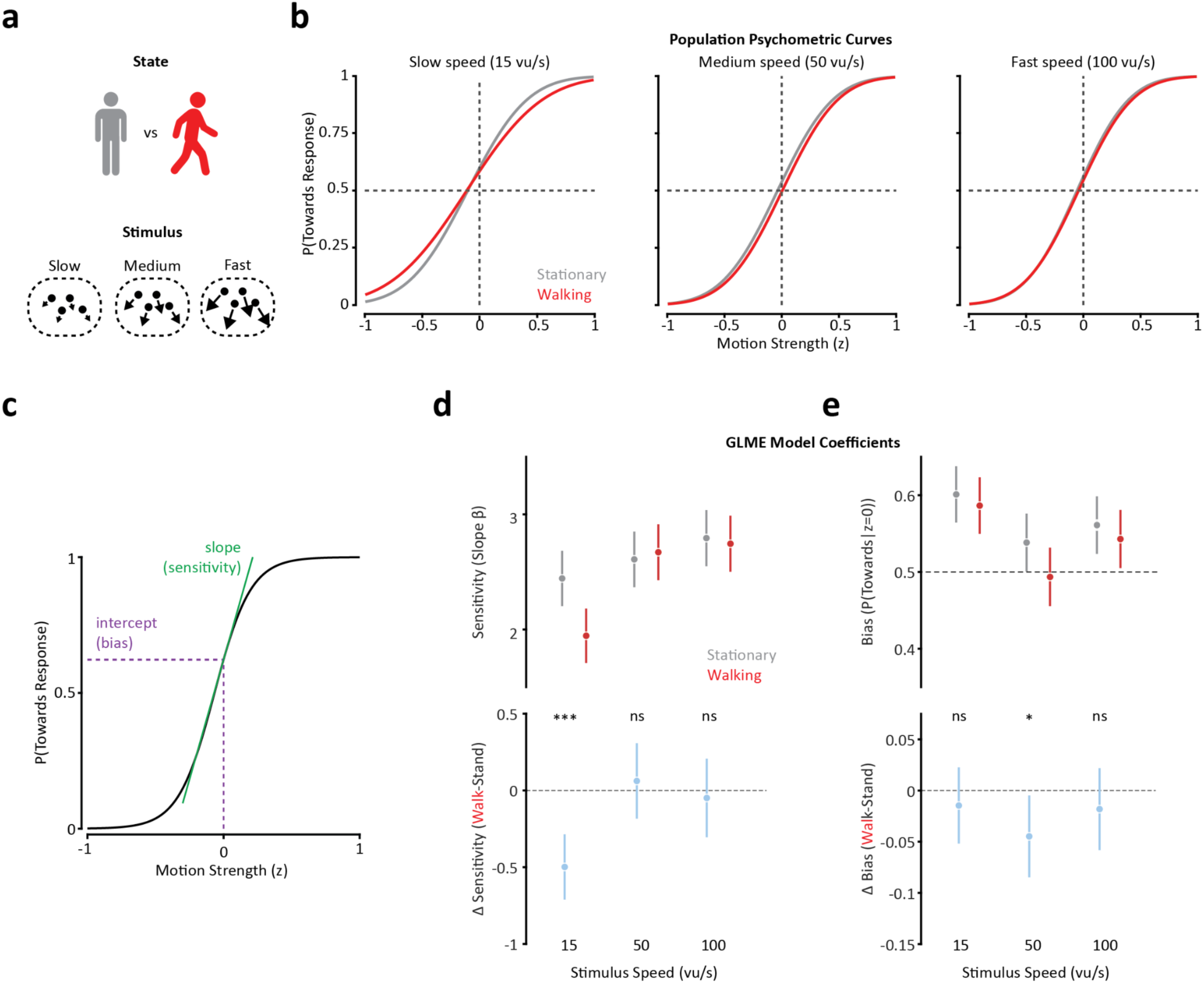
Locomotion dissociably alters perceptual sensitivity and bias, dependent on optic flow speed. a) Overview of the experimental conditions. We presented trials with one of three different stimulus speeds while participants were either stationary (grey) or walking (red). b) Population psychometric curves obtained from the Generalized Linear Mixed Effects (GLME) model for each experimental condition. Each stimulus speed is shown on a separate panel with curves for stationary (grey) and walking (red) trials overlaid. c) Schematic illustrating slope (sensitivity) and intercept (bias) coefficients obtained from the GLME model. d) Sensitivity was selectively reduced for slow optic flow during walking. *Top panel*: Standardized regression coefficients (*β*) for the slope parameter. Markers indicate the point estimate and error bars represent standard error of the mean. *Bottom panel*: Pairwise differences between the coefficients (Walking-Stationary). The solid line at Δ = 0 represents the null hypothesis of no difference in predictive weight. Error bars for the differences represent 95% confidence intervals. e) Walking promoted an ‘away’ bias selectively for the medium stimulus speed. Conventions as in (d) for intercept coefficients converted to probabilities. *** p<0.001, * p<0.05, ns p>0.05; Pairwise *F*-tests on model coefficients.

Motion-in-depth sensitivity is selectively reduced for slow optic flow during locomotion. Independent of behavioral state, an ANOVA of the GLME model revealed that stimulus speed significantly modulated baseline sensitivity (F(2, 12056) = 4.05, p = 0.017), with sensitivity increasing at faster speeds (Figure 2d). This may be due to the increased effective motion strength provided by higher stimulus speeds, which scale the average motion vector. Evaluating the effects of locomotion revealed a significant three-way interaction between behavioral state, stimulus speed, and motion strength (*F*(2, 12056) = 6.65, *p* = 0.001), demonstrating that walking does not uniformly alter task performance, but instead that locomotion-induced changes in sensitivity are dependent on stimulus speed. To decompose this interaction, we examined planned condition-specific changes in psychometric slopes to determine how behavioral state modulated sensitivity across each stimulus speed. For the slow optic flow condition, the population psychometric curve exhibited a decreased slope during walking (Figure 2b), reflected by a significantly lower slope model weight compared to the stationary state (Figure 2d; *F*(1, 12056) = 21.1, *p* < 10^−7^). By contrast, at medium and fast speeds, we observed no significant group-level change in sensitivity during locomotion. Thus, locomotion resulted in a selective deficit for slow motion-in-depth sensitivity while leaving sensitivity to faster speeds intact.

Motion-in-depth bias was selectively modulated during walking for medium-speed optic flow. Across both behavioral states, participants exhibited a baseline ‘towards’ bias - a tendency to perceive ambiguous stimuli as approaching rather than receding. This baseline bias followed a non-monotonic function of stimulus speed (*F*(2, 12056) = 5.28, p = 0.005), being strongest at slow speeds and weakest at the medium speed (Figure 2e). While an omnibus ANOVA revealed no significant overarching interaction between behavioral state and stimulus speed for perceptual bias (p = 0.52), planned contrasts between behavioural states for each stimulus speed indicated that locomotion may selectively promote a significant relative ‘away’ bias at the medium stimulus speed (*F*(1, 12056) = 4.83, p = 0.028). Notably, this was the only experimental condition in which the population-level bias shifted to perceiving ambiguous motion as moving ‘away’. In light of the non-significant omnibus interaction, we treat this speed-specific bias shift as preliminary. Crucially, we investigate below whether this statistical ambiguity in the overarching effect of walking is resolved by moving beyond a simple binary behavioural state classification to evaluate how optic flow perception continuously covaries with the kinematics of locomotion gait.

Taken together, our results demonstrate that locomotion significantly modulates the perception of optic flow motion-in-depth. Confirming our GLME model, we found similar results using a two-stage analysis approach: fitting psychometric functions individually for each participant and statistically comparing the resultant parameters (Supplementary Figure 3). Alongside equivalent changes in sensitivity and bias, this individual-participant analysis revealed a significant increase in estimated lapse rates for slow optic flow during locomotion (Supplementary Figure 3d). This indicates that participants found the slow stimulus speed condition particularly challenging during walking, in line with previous findings investigating visual speed estimation ^17^. Importantly, we found that locomotion does not simply induce a global change in task performance. Instead, sensitivity and bias are dissociably dependent on optic flow speed. These speed-dependent effects provide strong evidence against a generic behavioral state change such as generalized arousal or cognitive load, but instead are evidence that specific sensorimotor mechanisms drive changes in optic flow perception during walking.

### Step frequency dynamically modulates perceptual sensitivity and bias for optic flow motion-in-depth

To determine whether the modulation of optic flow perception during walking reflects a binary behavioral state change or is continuously tuned by gait kinematics, we evaluated how task performance covaries with trial-by-trial variations in step frequency, a reliable cue for self-motion speed estimation^37^. We determined step frequency from the rhythmic vertical oscillations of the head over the stepping cycle (Figure 1a,c; Figure 3a). We observed substantial natural intra-participant variance in step frequency across trials (mean ± SEM step frequency averaged over the duration of each trial = 1.54 ± 0.04 Hz, corresponding to 0.72 ± 0.03 m/s walking speed; mean intra-participant range: 0.47 ± 0.02 Hz), providing a natural continuous variable to test for covariation with task performance. Linear mixed-effects modeling confirmed that stimulus speed did not influence step frequency, likely owing to the short trial durations used (0.6 s). To account for baseline biomechanical differences between individuals, step frequency was z-scored within each participant.

**Figure 3:**
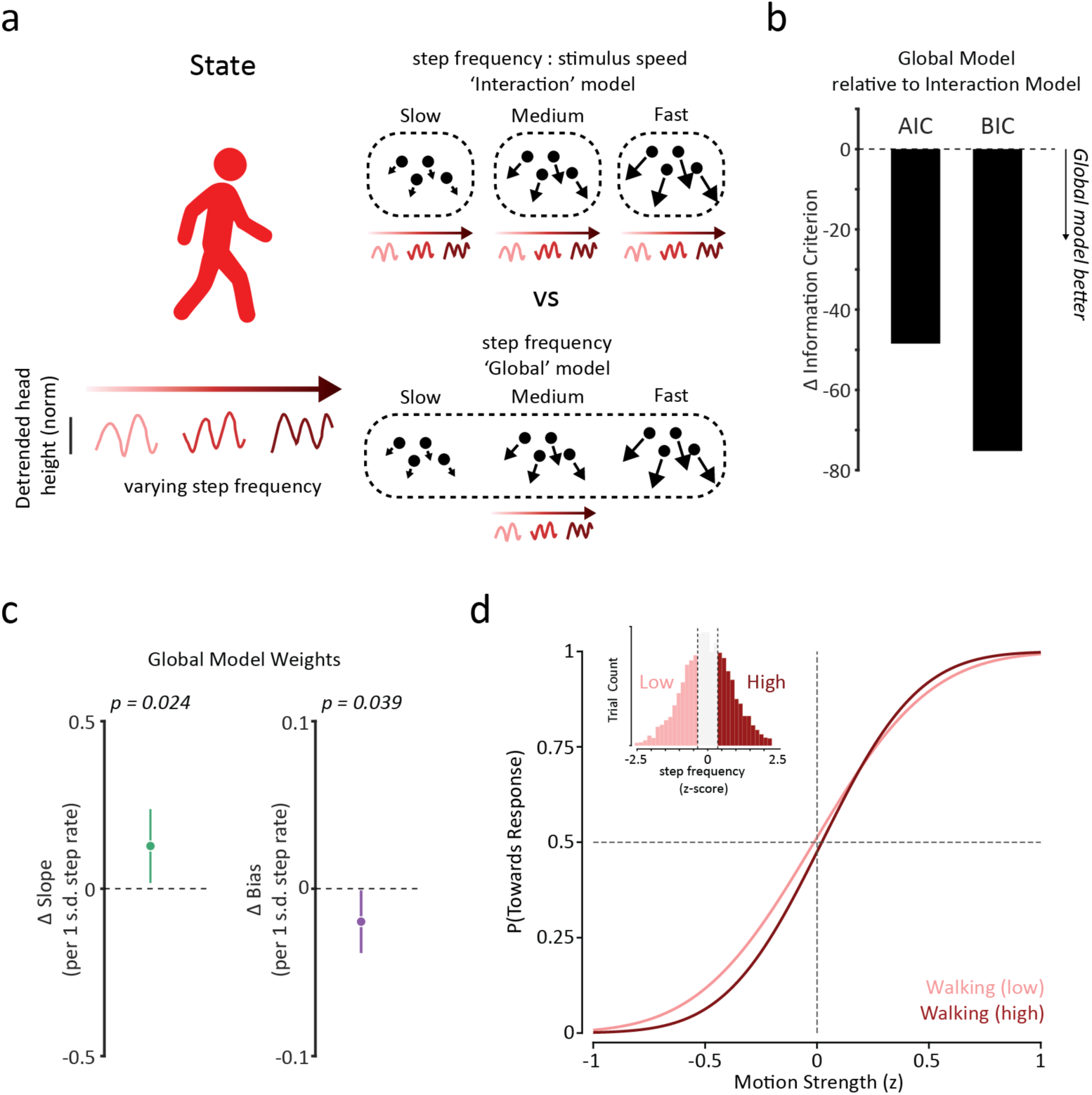
Perceptual sensitivity and bias covary with step frequency. a) We investigated how step frequency alters optic flow perception by comparing an ‘interaction’ model, in which stimulus speed-specific model weights were obtained for the effects of step frequency, and a ‘global’ model, in which global model coefficients were obtained for the effects of step frequency independent of stimulus speed. b) Akaike and Bayesian Information Criteria both indicated that the global step frequency model provided a more parsimonious fit to the data. c) Model weights for the effects of step frequency on sensitivity (slope) and bias (intercept). Increasing step frequency is associated with increased perceptual sensitivity and an ‘away’ perceptual bias. d) Population psychometric curves for low and high step frequency walking. We partitioned walking trials based on step frequency tertiles.

The modulatory effect of step frequency is invariant to optic flow speed. Because the difference in task performance between stationary and walking states was stimulus speed-specific, we tested whether the influence of step frequency similarly varied across stimulus speeds. We fit a GLME model to walking trials in which step frequency could independently influence bias and sensitivity for each stimulus speed (Figure 3a; ‘interaction’ model). An ANOVA of this model revealed no significant interaction between step frequency and stimulus speed for either perceptual sensitivity (p=0.78) or bias (p=0.43). Indeed, the stimulus speed-specific model weights trended in the same direction across all three stimulus speeds (Supplementary Figure 4). We therefore tested whether a simpler model in which single global step frequency weights applied to all stimulus speeds fit the data better (Figure 3a; one weight for intercept, one for slope). This ‘global’ model yielded superior goodness-of-fit metrics compared to the ‘interaction’ model (ΔAIC = 48.46; ΔBIC = 75.21; Figure 3b). This suggests that while the discrete behavioral state change between being stationary and walking depends on optic flow speed, the continuous modulation of optic flow perception with step frequency is largely independent of optic flow speed.

Perceptual sensitivity and bias for optic flow covary with step frequency. Having established the global relationship between step frequency and stimulus speed, we examined the fixed-effect coefficients of the model to determine how step frequency alters optic flow perception. We found that higher step frequencies were associated with a significant increase in perceptual sensitivity (Figure 3c; *β*_slope_ = 0.13 ± 0.06 per standard deviation of step frequency, *p* = 0.024). Simultaneously, increasing step frequency promoted a progressively stronger relative bias to perceive optic flow as moving ‘away’ (Figure 3c; *β* _bias_ = −0.05 ± 0.02, *p* = 0.039) To visualize these continuous effects, we partitioned walking trials into tertiles based on step frequency (Figure 3a,d). Consistent with the continuous model, comparing the lowest and highest tertiles revealed that high step frequency walking both steepens the psychometric slope and shifts the curve rightward (in the direction of an ‘away’ bias) compared to low step frequency walking (Figure 3d).

### High step-frequency walking rescues sensitivity deficits and promotes an ‘away’ bias compared to the stationary state

Differences in optic flow perception between stationary and walking states depend on step frequency. We next directly compared these gait-dependent changes in task performance back to the stationary baseline. We first fit a discrete GLME model pooling trials across all stimulus speeds to compare stationary trials against the low and high step frequency walking tertiles (Figure 4a). Walking at a low step frequency resulted in a significant global sensitivity deficit compared to the stationary baseline (Figure 4b; Δ*β* = −0.27 ± 0.13, F(1, 7964) = 4.38, p = 0.036). However, walking at a high step frequency completely reversed this deficit, restoring perceptual sensitivity to stationary baseline levels (Figure 4b, d; Δ*β* = +0.23 ± 0.14, F(1, 7964) = 2.69, p = 0.101; comparison between high and low step frequency: Δ*β* = +0.49 ± 0.16, F(1, 7964) = 8.99, p = 0.0027). Simultaneously, comparing perceptual bias revealed that while low step frequency walking did not significantly alter directional bias (F(1, 7964) = 0.99, p = 0.319), high step frequency walking induced a robust and significant relative ‘away’ bias compared to the stationary baseline (Figure 4c, d; ΔZ = −0.15 ± 0.06, F(1, 7964) = 6.82, p = 0.0090). This dynamic gait-dependency clarifies the statistical ambiguity of our initial binary state analysis (Figure 2e), demonstrating that the shift in bias during walking is contingent on the kinematics of walking rather than just a binary state change.

**Figure 4:**
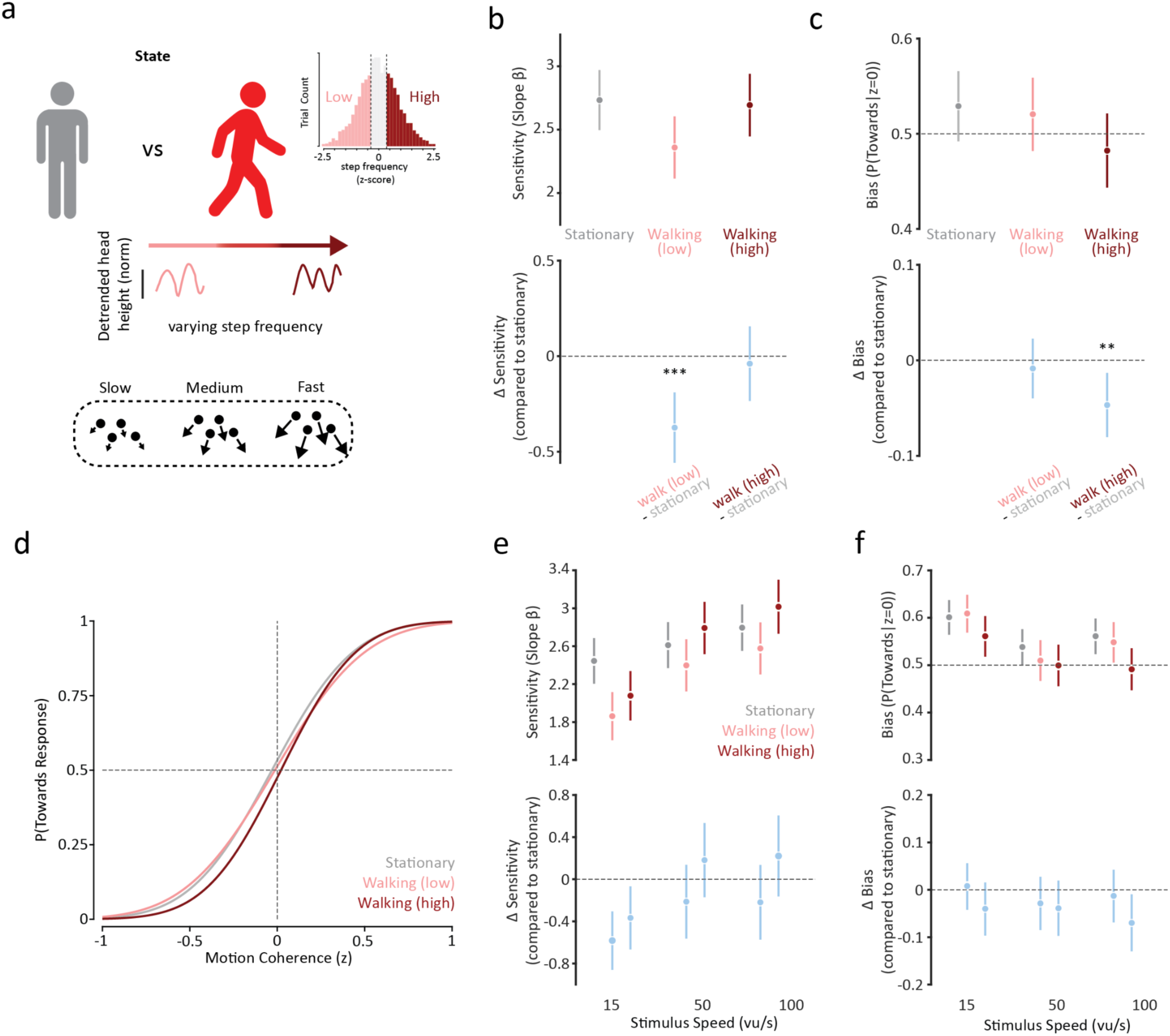
High step frequency walking rescues visual sensitivity and promotes an ‘away’ perceptual bias. a) We compared task performance between stationary and low and high step frequency walking, pooling trials across stimulus speeds. b) T*op panel*: Standardized regression coefficients (*β*) for the slope parameter for the Stationary, Walking (low step frequency) and Walking (high step frequency) states. Markers indicate the point estimate and error bars represent standard error of the mean. *Bottom panel*: Pairwise differences between the stationary baseline and low and high step frequency walking (Walking low/high - Stationary). The solid line at Δ = 0 represents the null hypothesis of no difference in predictive weight. Error bars for the differences represent 95% confidence intervals. c) Same as (b) for bias. d) Population psychometric curves for stationary (grey), low step frequency walking (pink) and high step frequency walking (red), for the ‘Global’ step frequency model. e) Same as (b) for the speed:step frequency interaction model. f) Same as (c) for the speed:step frequency interaction model. *** p<0.001, * p<0.05, ns p>0.05; Pairwise *F*-tests on model coefficients.

Given that the binary effects of walking were stimulus speed-dependent (Figure 2d, e), we next asked how varying step frequency shaped task performance relative to the stationary baseline across optic flow speeds (Figure 4e, f). For visual sensitivity, high step frequency walking did not fully rescue the deficit for slow optic flow back to baseline levels, but it fully restored sensitivity for medium and fast optic flow, even exceeding the stationary baseline (Figure 4e). In parallel, high step frequency walking produced a consistent ‘away’ shift in perceptual bias relative to the stationary baseline across all stimulus speeds (Figure 4f). For slow and fast stimulus speeds, this ‘away’ bias was markedly stronger during high compared to low step frequency walking, whereas for medium speeds, both stepping tertiles showed an equally pronounced shift away from baseline. Ultimately, these kinematic-dependent modulations demonstrate that visual perception is dynamically calibrated by ongoing gait: high step frequency both restores sensitivity to behaviorally relevant optic flow speeds and progressively biases judgments toward ‘away’ motion, consistent with the active cancellation of expected self-generated flow.

## Discussion

How does behaviour shape sensory perception? We developed a novel 3D motion coherence paradigm to investigate this question in the context of how locomotion shapes optic flow motion-in-depth perception. One possibility is that locomotion results in a general degradation of perception due to increased sensory noise and reduced available cognitive resources ^38–41^. Alternatively, the brain may automatically leverage sensorimotor contingencies to actively modulate optic flow perception^25,26^. Our findings provide strong evidence for the latter. We found that walking drives dissociable changes in perceptual sensitivity and bias for optic flow motion-in-depth, dependent on optic flow speed. These modulations by walking are not a binary response to the onset of movement. Instead, both sensitivity and bias scale continuously with step frequency - a reliable idiothetic cue for self-motion speed ^37,42^. Taken together, our findings suggest that the visual system adapts to both the expected visual statistics of locomotion and leverages real-time non-visual motor signals to dynamically calibrate optic flow perception.

Locomotion induces a relative shift in perceptual sensitivity toward faster optic flow speeds. Sensitivity to slow optic flow was selectively impaired during walking while performance at faster speeds was preserved. This speed-dependent profile converges with findings from a distinct visual speed discrimination task in which locomotion similarly degraded performance at slow but not fast optic flow speeds ^17^. The emergence of this pattern across both coherent speed discrimination and stochastic 3D motion-in-depth direction discrimination argues against task-specific cognitive strategies and instead indicates a more general upstream sensory mechanism that shifts the operating range of visual motion processing toward faster, more behaviourally relevant speeds during locomotion ^8,43^. Our results therefore cannot be explained solely by an increase in sensory noise and cognitive load during walking. Indeed, during high step frequency walking, perceptual sensitivity for medium and fast optic flow trended higher than stationary baseline levels, despite the heightened cognitive and motor demands of walking while performing the task wearing a head-mounted display. This relative shift in visual sensitivity aligns with an efficient coding framework, wherein the visual system dynamically reallocates limited sensory resources to match the higher velocity statistics inherent to locomotion ^17,44,45^.

Locomotion systematically recalibrates directional bias, driving an active cancellation of predicted self-generated optic flow. During stationary trials, participants exhibited a baseline bias to perceive ambiguous stimuli as approaching ^46^. Under a naive Bayesian framework, forward locomotion should establish an expectation for expanding optic flow, pulling ambiguous percepts toward that prior and increasing ‘towards’ reports. Instead, walking produced the opposite effect: the baseline towards bias was attenuated and, in some cases, actively reversed into a small net ‘away’ bias. This repulsive shift is consistent with an internal subtractive flow-parsing mechanism, in which expected self-motion is subtracted from incoming retinal flow to isolate external motion in depth ^32^, converging with prior reports that non-visual walking signals discount perceived visual speed ^16,17,27,29,30^. Previous investigations demonstrating non-visual contributions to flow-parsing relied on closed-loop virtual environments with textured ground planes and visual landmarks ^47^. Crucially, because our paradigm utilized a strictly open-loop display without a ground plane or visual references, our findings demonstrate that non-visual motor signals alone are sufficient to drive subtractive flow-parsing in depth, in the complete absence of closed-loop visual feedback.

These perceptual modulations are not a fixed, binary property of the walking state. By tracking trial-by-trial gait, we found that both sensitivity and bias scale continuously with step frequency^37,42^. Walking at a low step frequency produced a significant sensitivity deficit compared to the stationary trials, whereas high step frequency walking completely rescued this deficit, restoring performance to stationary baseline levels. Simultaneously, high step frequency walking drove a robust and significant ‘away’ bias relative to the stationary baseline. Model comparisons revealed that step frequency scaled both sensitivity and bias globally across all stimulus speeds, with a speed-invariant model significantly outperforming a speed-interaction model. Thus, visual processing does not simply toggle into a fixed ‘locomotion mode’ upon movement onset; rather, the brain continuously leverages real-time idiothetic cues to dynamically tune optic flow perception to the ongoing rhythm of gait.

Are changes in perceptual sensitivity and directional bias governed by separate mechanisms, or do they reflect a unified process? Distinct speed profiles between stationary and walking trials superficially suggest separate operations - sensory noise filtering at slow speeds and predictive subtraction at medium speeds. However, both phenomena can be reconciled within the framework of efficient sensory coding ^44,45^. Durgin & Gigone^17^ proposed that concurrent changes in sensitivity and bias could arise from adaptive decorrelation implemented via subtractive inhibition, a model based on Barlow’s work ^48,49^. The subtractive inhibition of visual speed perception drives the shift in perceptual bias. The decorrelation of visual and non-visual self-motion signals recalibrates coding dynamic range to modulate visual sensitivity. Our findings could also be explained by a Bayesian model of efficient coding ^50^. Under this framework, sensory resources (quantified using Fisher information) are dynamically allocated to match environmental priors: during locomotion, the visual system reallocates finite coding capacity to encode motion aligned with forward travel - specifically faster, expanding optic flow. From this framework, two corollaries follow. First, concentrating resources on locomotion-congruent speeds reduces resources for encoding slower speeds, explaining our selective loss in sensitivity. Second, Bayesian decoding over this asymmetric representation induces a skewed likelihood function that mathematically repels perceived directions away from the region of highest coding density. Thus, dedicating more resources to encode expanding ‘towards’ motion can paradoxically cause a relative ‘away’ bias, as we found here. Crucially, our kinematic analysis demonstrates that step frequency simultaneously scaled both sensitivity and bias across all speeds, providing behavioral evidence that an idiothetic signal recalibrates both dimensions in unison.

Our psychophysical findings predict that locomotion-dependent changes in neural speed tuning may be conserved across mammalian species. In mice, locomotion shifts visual cortex tuning toward faster speeds^43^, consistent with the relative reallocation of coding resources towards faster optic flow speeds that we found. Such shifts could also produce our observed perceptual biases through an alternative, non-normative mechanism: if downstream decoders remain fixed to stationary-state tuning rather than updating to changes in neural tuning during locomotion, the readout will misattribute neural activity to slower visual inputs ^8^, systematically underestimating optic flow speed, consistent with the known subtractive effect of walking on perceived visual speed^16,17,27,29,30^. If this tuning shift occurs asymmetrically stronger in neurons selective for congruent, expanding optic flow (as has been found perceptually in humans ^16^), an uncorrected readout would cause the relative “away” bias we observed. Primate optic flow-encoding areas such as MST are modulated by non-visual self-motion signals ^24^, and key principles of locomotion modulation are conserved between mouse V1 and peripheral primate V1^19^, making parallel tuning shifts in primates plausible. Within this framework, our findings predict that step frequency acts as a continuous physiological calibration signal, dynamically scaling both the magnitude of cortical tuning shifts and the precision of visual motion representations.

### Limitations of the study

To isolate non-visual self-motion signals, our paradigm intentionally simplified several features of natural navigation. An open-loop rendering design was essential to eliminate visual motion confounds from head movement, ensuring that any perceptual modulation is attributable to idiothetic cues rather than differences in retinal input. Similarly, the absence of stable background objects or a ground plane ensured that participants relied strictly on the optic flow stimulus rather than external visual reference frames ^51–53^. While our discrete 2-AFC design and brief walking intervals enabled rigorous psychophysical modeling, they necessarily simplify the continuous, graded decisions and adaptation dynamics of prolonged locomotion ^35,54^. Furthermore, the resulting visuomotor mismatch and added cognitive load of navigating with a head-mounted display likely increased the task difficulty during walking and may have dampened our observed effects of walking ^55^. These deliberate constraints nevertheless enabled the clean attribution of perceptual changes to non-visual locomotion signals. Future closed-loop paradigms and continuous psychophysical methods ^56,57^ can build on this foundation to test how these mechanisms generalise to extended natural navigation.

### Conclusions

Our findings demonstrate that human optic flow perception is actively and dynamically tuned by ongoing locomotion. By revealing that visual sensitivity and bias covary with step frequency, we show that the brain utilizes the body’s physical rhythm as a calibration signal to resolve sensory ambiguities. This dynamic scaling ensures visual processing matches the demands of physical translation, underscoring that human vision is fundamentally optimized for the active, moving observer.

## Methods

### Participants

Sixteen healthy volunteers (10 female, 6 male; 12 aged 19–28 years old, 2 aged 29-38 years old, 2 aged 39–48 years old) completed the task using the same equipment and procedures. All participants had normal or corrected-to-normal vision. Where vision was corrected contact lenses were used to allow the comfortable wearing of the head-mounted display. All participants were recruited from the UCL SONA experimental participant database in exchange for payment. All participants provided informed written consent before participating in the experiment. All experiments were approved by the Ethics Committee of the Department of Experimental Psychology, UCL (EP/2019/002).

### Visual Stimulus and Task

#### Apparatus and Experimental Setup

Stimuli were generated using Unity (Unity Software Inc.) presented stereoscopically using a Vive Pro Eye head-mounted display (HMD; HTC Corporation) with an approximate horizontal field-of-view of 100° and 90 Hz refresh rate. The HMD was interfaced with the host computer using a wireless adapter. Trial presentation and data collection were controlled using custom-written scripts in MATLAB (MathWorks). Global and trial-specific parameters were sent from MATLAB to Unity, and response data from Unity to MATLAB, using a TCP-IP protocol.

For the locomotion conditions, participants walked along a straight path defined by a guide rope (∼8m) which they were loosely attached to via a carabiner and lightweight climbing belt (Supplementary Figure 1). Subjects were instructed to briefly touch the rope at the start of a block of trials to ensure correct allocentric orientation within the room, but to refrain from touching it during trials.

#### Optic Flow Motion Coherence Stimuli

The experiment consisted of a single-interval (duration = 0.6 s), two-alternative forced-choice (2AFC), motion-in-depth direction discrimination task. Throughout all trials and within-block inter-trial intervals, participants were presented with a mid-grey background containing a centrally located black fixation cross. Stimulus kinematics and dimensions were defined using standard Unity virtual units (vu; see Supplementary Figure 2 for related visual speeds).

The optic flow stimulus consisted of 175 white spheres (diameter = 2 vu). At the beginning of each trial, spheres were distributed at random 3D coordinates within an invisible 100 vu diameter spherical stimulus bounding space (Figure 1; Supplementary Figure 2). Each sphere was assigned a constant 3D velocity vector along the azimuth plane (with fixed elevation), randomly sampled from a Von Mises distribution. This distribution was parameterized by a mean direction (μ) constrained to μ = 0° for ‘towards’ trials or μ = 180° for ‘away’ trials, and a concentration parameter (κ) ranging from 0.01 to 100 which determines the circular variance of the distribution. The magnitude of each sphere’s velocity was scaled to the trial speed of 15 vu/s (slow), 50 vu/s (medium) or 100 vu/s (fast). In this context, κ acts as a manipulation of the signal-to-noise ratio governing motion strength. A high κ value (e.g., 100) generates a highly concentrated distribution where nearly all spheres travel in the mean direction (Figure 1b). Conversely, a low κ value (e.g., 0.01) flattens into a nearly uniform circular distribution (high noise).

To maintain uniform density and prevent synchronous respawning artifacts, spheres were initialized with a random starting age assigned uniformly between 0 and 0.2 s, and had a limited lifetime of 0.2 s, after which they were randomly teleported to a new coordinate within the volume. Additionally, if a sphere’s trajectory carried it beyond the geometric bounds of the spherical stimulus volume it was also randomly teleported to a new coordinate within the sphere.

To ensure approximately uniform coverage of the visual field without unnatural edge artifacts, the virtual camera was positioned 70 vu offset along the depth axis from the center of the spherical stimulus bounding volume. Subjects therefore observed the stimulus from an external vantage point, with an effective near clipping frame of 20 vu. The camera was also configured with a far clipping plane of 60 vu. A spherical bounding volume was chosen over a cubic volume to ensure rotational invariance and prevent abrupt visual transients. Because the boundary is uniformly curved, the spatial profile of the stimulus remains identical regardless of the global motion direction. This ensures that direction discrimination is driven purely by optic flow signals, rather than the sudden appearance or disappearance of sharp geometric corners formed by the abrupt teleportation of spheres.

Notably, the virtual environment contained no pictorial cues for absolute scale. The sole anchor for metric depth was the physical and simulated interpupillary distance (IPD), which was optimized for each participant using the native calibration routine of the Vive Pro Eye. Given the abstract nature of the environment, this binocularity primarily provided a geometric reference for relative depth, not absolute scale or speed. Furthermore, visual rendering was strictly open-loop; the position and rotation of the virtual cameras remained fixed relative to the global stimulus trajectory and did not update based on the participant’s physical head movements.

### Experimental Procedure

Subjects were tested across 6 randomly interleaved experimental conditions varying across 2 factors: behavioral state (stationary vs. walking) and stimulus speed (15, 50, and 100 vu/s).

Before data collection, participants practiced walking with the HMD and guide rope. They also completed a set of practice trials (κ = 1, 10, 50) with visual feedback (a green/red circle around the fixation cross for correct/incorrect responses). During the main experiment, no feedback was provided.

The experiment was structured into blocks of 4 trials, consisting of 2 stationary trials followed by 2 walking trials, or vice versa. Each 4-trial block was completed during a single traversal of the physical experimental space. The stimulus speeds on each trial were randomly interleaved.

Each trial lasted 0.6 s, ending with the sudden disappearance of the spheres. Subjects then verbally reported their perceived direction of global motion (‘towards’ or ‘away’) which was entered into the computer by an experimenter who was blind to the stimulus shown. Subjects were offered an optional pause every 40 trials and were required to take a 5-minute break without the HMD every 160 trials.

### Adaptive Psychometric Trial Presentation

Within each of the 6 experimental conditions, stimulus difficulty was controlled by varying the Von Mises concentration parameter (κ) using a combination of constant stimuli and 3-down, 1-up staircases (targeting ∼79% convergence). Constant stimuli consisted of 5 repeats of specific κ values (0.025, 0.4, 0.8, 1.4, 2.5, 5) for both ‘towards’ and ‘away’ directions. Staircases utilised a broader pool of κ values (0.01 0.025 0.05 0.1 0.2 0.4 0.8 1 1.2 2.5 5 10 25 50 75 100). Four independent 18-trial staircases were run per condition: two for the ‘towards’ direction and two for the ‘away’ direction, starting at either an easy (κ = 5) or hard (κ = 0.1) difficulty level. In total, participants completed 132 trials per experimental condition, yielding 792 total trials across the entire experiment.

### Trial inclusion

From the total scheduled dataset of 12,672 trials (16 participants were filtered through a sequential rejection protocol: × 792 scheduled trials), data

1. *Technical and Response Gating:* 406 trials (3.20%) were excluded due to corrupted data packets, experimenter input timeouts/invalid response codes, or wireless head tracking dropouts during the 0.6 s stimulus presentation window, leaving 12,266 recorded trials.
2. *Step Frequency Outlier Removal:* To eliminate stepping pauses and gait tracking outliers, walking trials falling in the extreme 0.5% upper and lower tails of trial-averaged step frequency were trimmed on a within-participant basis, removing an additional 198 trials (1.61% of recorded trials).

In total, 604 trials (4.77% of scheduled trials) were excluded across the entire study, leaving 12,068 valid trials across the 16 observers for all mixed-effects psychometric analyses. Data retention was consistent across participants (mean participant retention: 95. 2% ± 2. 1%; 12 of 16 observers retaining > 98% of scheduled trials).

### Psychometric Modeling and Statistical Analysis

To quantify perceptual performance, the 3D directional vectors of the optic flow stimulus (sampled from a Von Mises distribution) were first transformed into a continuous 1D scalar of signed motion strength (StimulusStrength). This value was defined as the mean cosine projection of all individual sphere trajectories onto the depth (z) axis, where positive values represented net motion ‘towards’ the observer and negative values represented net ‘away’ motion:

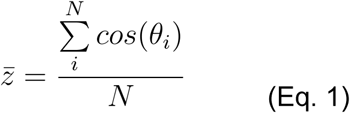

where *N* is the total number of spheres indexed by the subscript *i* and *θ_i_* is the angle between the ‘towards’ direction and the motion vector of sphere *i*.

Population-level psychophysical data were analyzed using Generalized Linear Mixed-Effects (GLME) models, implemented via the *fitglme* function in MATLAB (MathWorks). All models utilized a Binomial response distribution and a Probit link function to relate the linear predictors to the probability of a ‘towards’ response. To account for inter-participant variability in baseline bias and sensitivity, models included a random-effects structure consisting of random intercepts and random slopes for motion strength grouped by Subject (1 + StimulusStrength | Subject).

Initially, a factorial GLME was employed to evaluate the overarching main effects and interactions of behavioral state (Stationary vs. Walking), stimulus speed (15, 50, and 100 vu/s), and motion strength

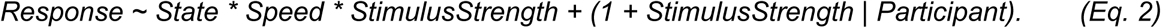

To derive absolute performance metrics for specific experimental conditions, the models were subsequently re-parameterized using dummy variable (cell-means) coding.

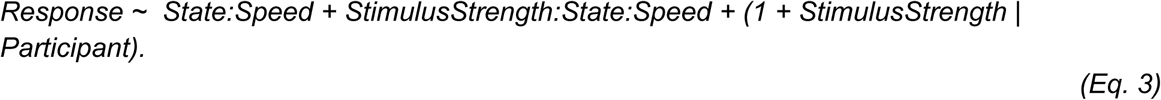

This approach allowed for the direct extraction of absolute Sensitivity (represented by the psychometric slope, i.e. interaction with *StimulusStrength*) and Bias (the intercept) estimates, along with their associated Standard Errors, from the model coefficient outputs. For visualization, intercept estimates (originally in probit units) were transformed into probabilities using the cumulative normal distribution function. Associated Standard Errors for these probability estimates were calculated using the Delta method (a first-order Taylor expansion) to account for the non-linear transformation.

Planned post-hoc comparisons were conducted to evaluate differences in these parameters across behavioral states using custom contrast vectors evaluated via Wald *F*-tests (Matlab function *coefTest*).

To investigate whether locomotion exerts a graded effect on task performance, we analyzed the continuous intra-participant variance in gait kinematics. Mean step frequency across trial duration (see below for estimation) was z-scored within participants to standardize relative variance across participants. We initially fitted a GLME to test for potential interaction effects between continuous step frequency and stimulus speed

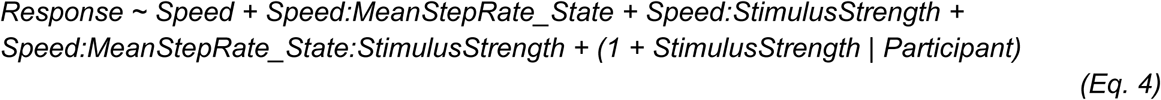

To determine if the influence of gait was speed-dependent, this full interaction model was compared against a more parsimonious ‘global’ model, which pooled the monotonic effects of step frequency across all stimulus speeds

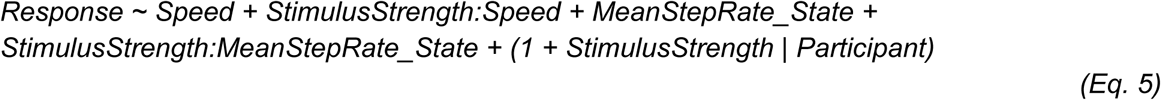

Model selection was guided by the Akaike Information Criterion (AIC) and Bayesian Information Criterion (BIC) which provided consistent findings.

To further visualize gait-dependent effects and facilitate direct comparison with the stationary baseline, walking trials were partitioned into tertiles (Low, Medium, and High step frequency). A discrete GLME was subsequently fitted to these categorical bins (*StepBin*) to extract condition-specific psychometric parameters

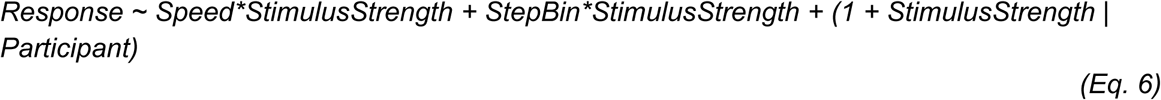

To compare walking with different step frequencies to stationary trials we created a behavioural state category (*StateCondition*) with categories *Stationary*, *Walking (low step frequency)*, and *Walking (high step frequency)*

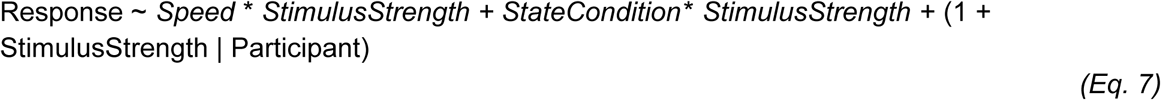

### Individual Psychometric Fits

To complement the population-level GLME models and explicitly quantify parameter changes at the level of individual participants, we fit separate psychometric functions for each participant across all combinations of behavioral state and stimulus speed. Psychometric curves were fitted using the Psignifit toolbox (Version 4; ^58^). For each condition, the probability of a ‘towards’ response was modeled using a cumulative normal distribution.

From these individual fits, three core metrics were extracted for subsequent statistical analysis. Sensitivity (Slope at Zero): The rate of change in response probability where motion strength (z) = 0. Bias (Intercept at Zero): The baseline propensity to report a ‘towards’ response where motion strength (z) = 0. Lapse Rate (λ): The fitted symmetric asymptote limit, representing the probability of a stimulus-independent response error.

Differences between the binary behavioral states (Stationary vs. Walking) were assessed using Wilcoxon Signed-Rank tests.

### Gait Analysis

To quantify the influence of locomotion kinematics on perceptual judgments, we extracted step frequency from continuous head-tracking data recorded from the HTC Vive Pro Eye. Initially, the continuous head-tracking data were segmented into discrete trial epochs. Each epoch was defined to begin 0.5 s prior to stimulus onset and conclude 0.5 s after the response was entered to capture the full movement profile during the decision-making process and enable sufficient sampling time to estimate step frequency.

During walking the head reaches its maximum vertical height during the mid-stance phase of a step, when the body’s center of mass passes directly over the supporting limb, and drops to its minimum height during the double-support phase following heel-strike. To isolate the rhythmic ‘head bob’ associated with walking, the continuous vertical position of the head (y-axis) was analyzed. This signal was first detrended using a moving average filter (1.0 s window). Step events were then identified by applying a peak-finding algorithm to the detrended vertical head position. A dynamic minimum peak prominence threshold was applied, defined as 0.5 × the standard deviation of the detrended y-position within that specific walking segment.

Instantaneous step frequency (Hz) was calculated as the inverse of the time interval between consecutive troughs. For each trial, the mean step frequency was calculated by averaging these instantaneous frequencies. To account for baseline biomechanical differences between individuals, these trial-averaged step frequencies were z-scored within each participant prior to psychometric modeling.

## Acknowledgements

We thank members of the Institute of Behavioural Neuroscience (IBN) for valuable discussion. This work was supported by a Biotechnology and Biological Sciences Research Council Doctoral Training scholarship to E.A.B.H.; a Laidlaw Undergraduate Research Scholarship to J.Y.L.T.; and the Sir Henry Dale Fellowship from the Wellcome Trust and Royal Society (200501), the Human Frontier in Science Program (RGY0076/2018), Bio-technology and Biological Sciences Research Council grant (BB/W01579X/1), and UKRI Frontier Research grant (EP/Y024656/1) to A.B.S.

## Supplementary Information

**Supplementary Figure 1:**
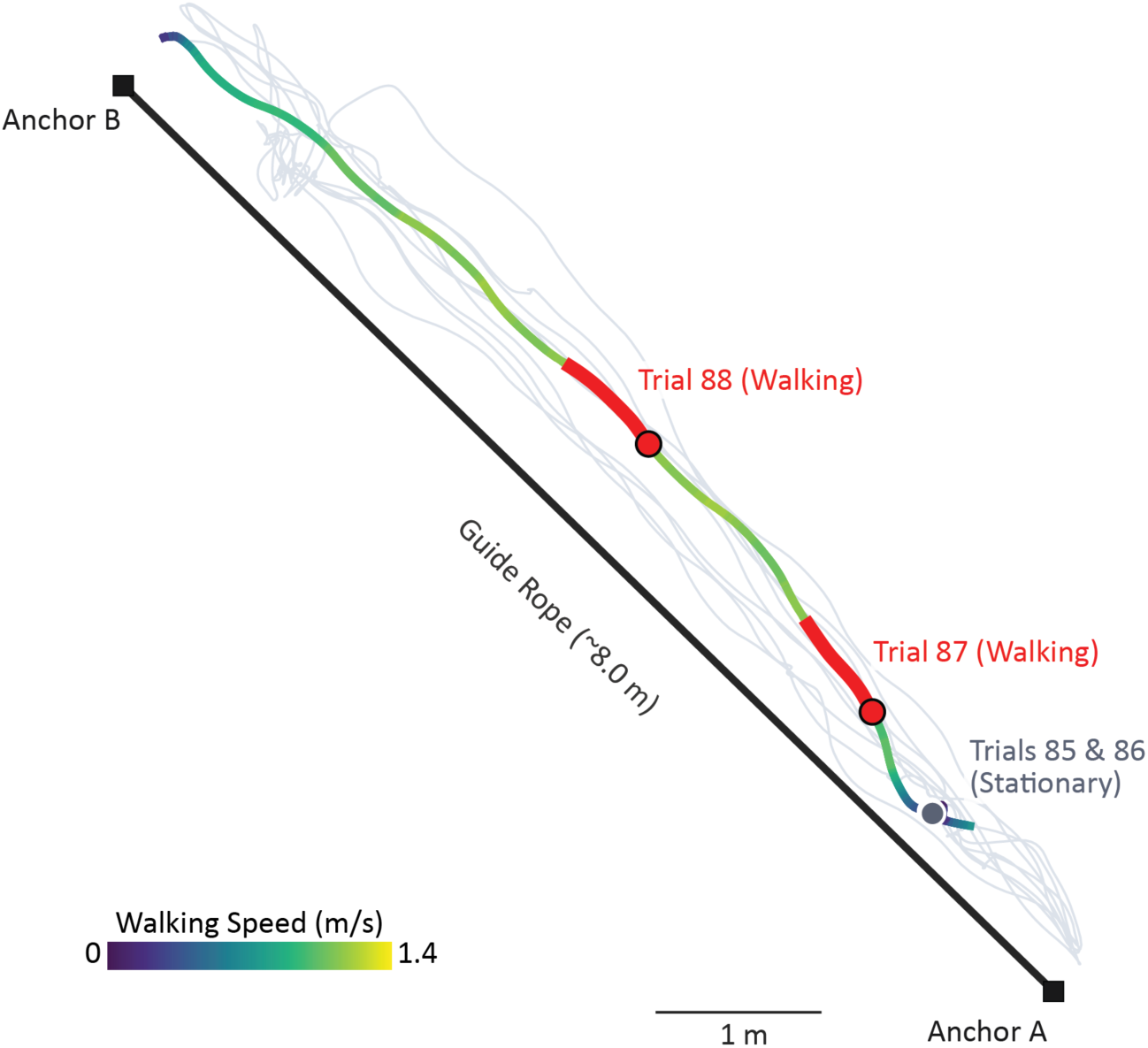
Top-down view of experimental area. Top-down schematic of the experimental area. Participants traversed an ∼8m straight path for each block of 4 trials. Example position trajectories are shown in grey for 10 blocks, with an active block colour-coded according to the participants walking speed. Grey circles indicate the location of stationary trials and red lines the location of walking trials in the active block. Participants were attached to a guide rope via a carabiner-linked belt around their waist.

**Supplementary Figure 2:**
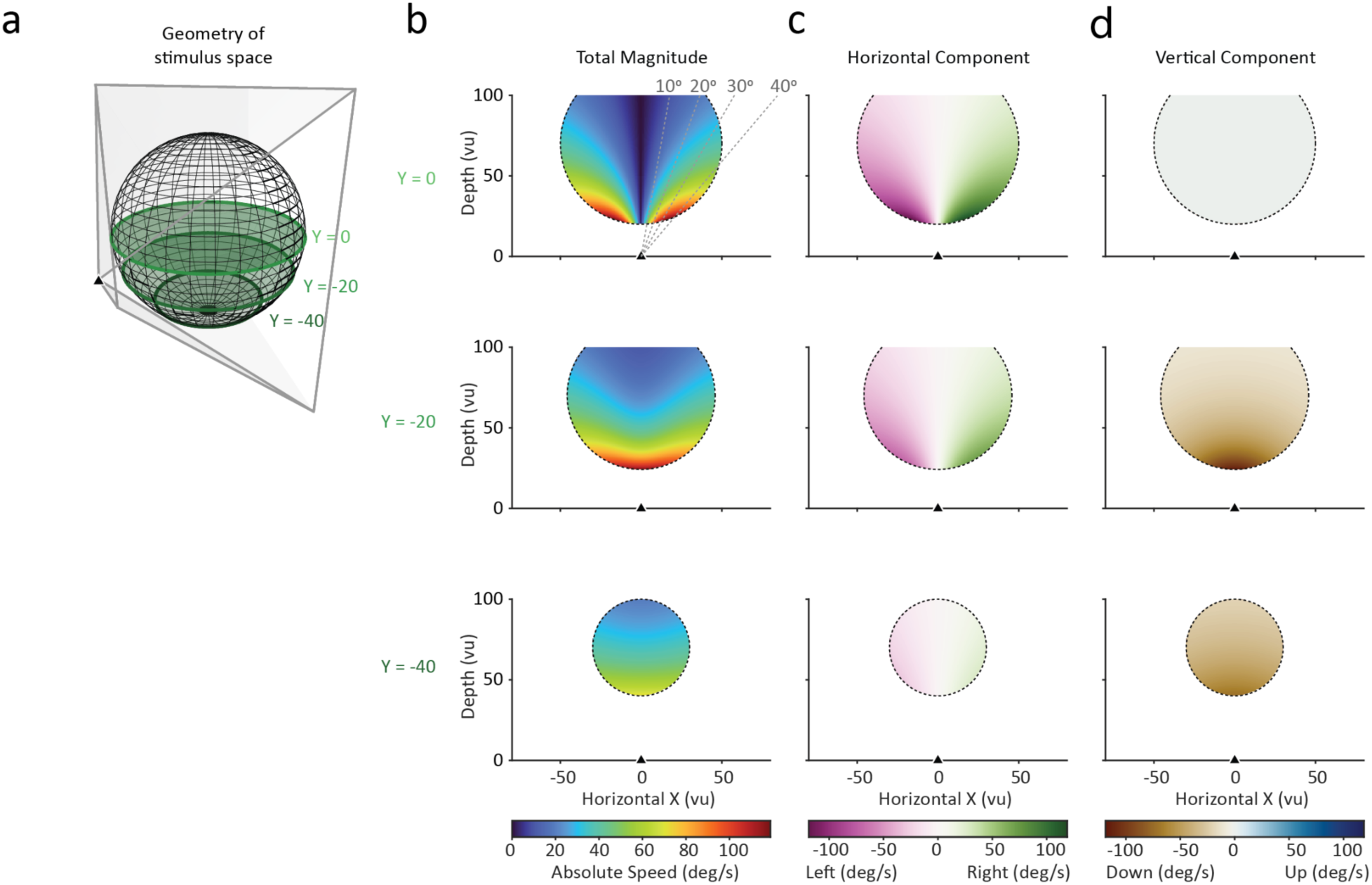
Geometry and visual speed profile of the stimulus. a) Schematic overview of the stimulus geometry. White spheres were spawned within a larger sphere volume (gridded outline shown). The observer was positioned outside the sphere volume (triangle) and viewed the stimulus through the camera frustum with near and far clipping frames (trapezium). Three different elevations are indicated by three green circular planes. b) Magnitude of visual speed (assuming stimulus speed of 100 vu/s and towards motion) in different parts of the geometric space. Note that the far clipping plane was at depth = 60 vu. c) Same as (b) for the horizontal component of visual speed d) Same as (c) for the vertical component of visual speed.

**Supplementary Figure 3:**
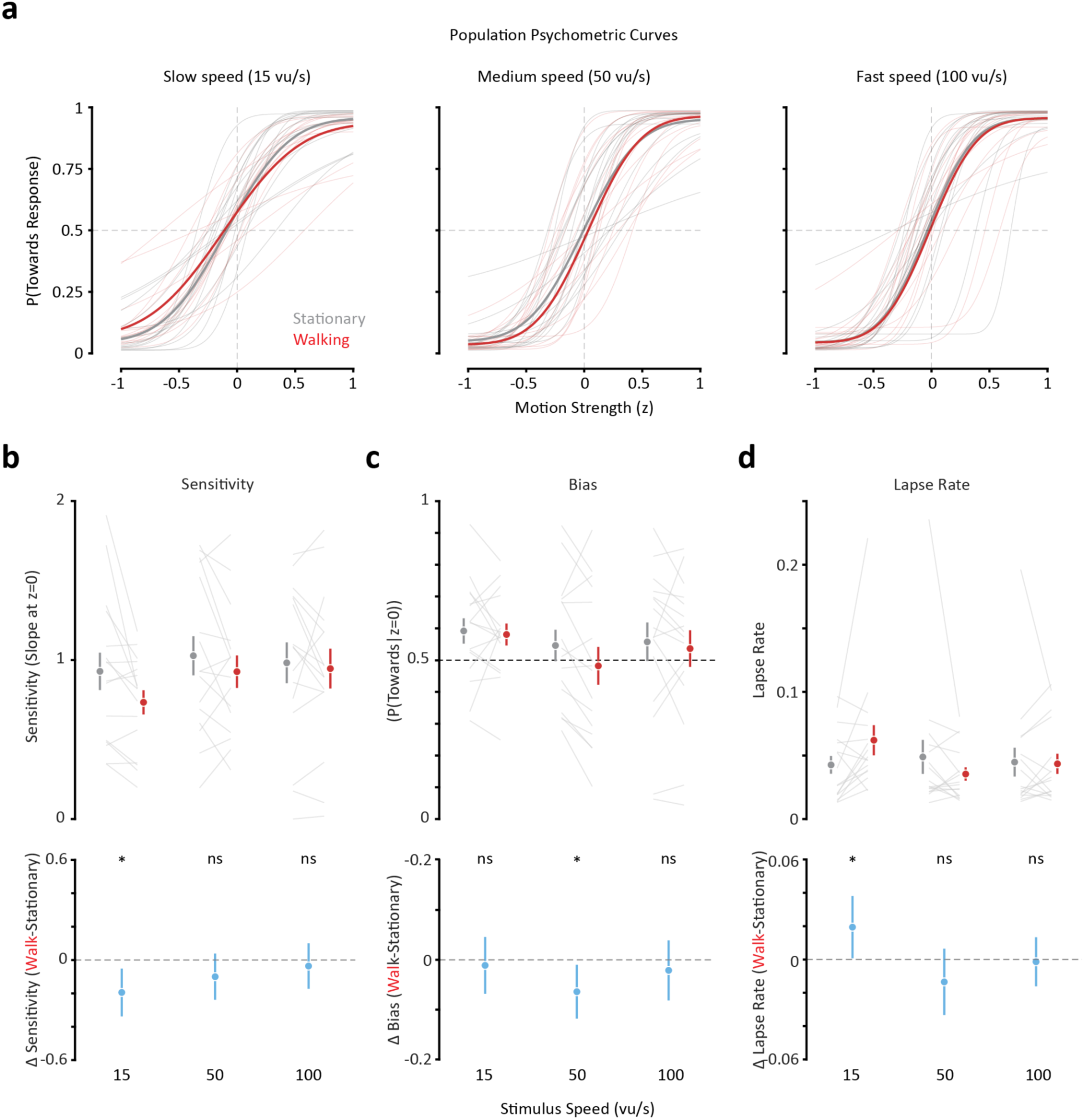
Two-stage psychometric function analysis replicates GLME findings. a) Population Psychometric Curves. Separate psychometric functions were fitted for each individual participant across all combinations of behavioral state (stationary (black) and walking (red)) and stimulus speed. Thick lines represent the population average curves, while thin lines represent individual participant fits. b) Sensitivity (Slope at z=0). The top panel shows individual (thin lines) and mean (markers) sensitivity for each speed, while the bottom panel shows the pairwise difference between walking and stationary states (mean ± 95% confidence intervals). c) Same as (b) for bias (P(Towards | z=0)). d) Same as (b) for lapse rate. Statistical significance for differences between behavioral states was assessed using Wilcoxon Signed-Rank tests (* p < 0.05, ns p > 0.05).

**Supplementary Figure 4:**
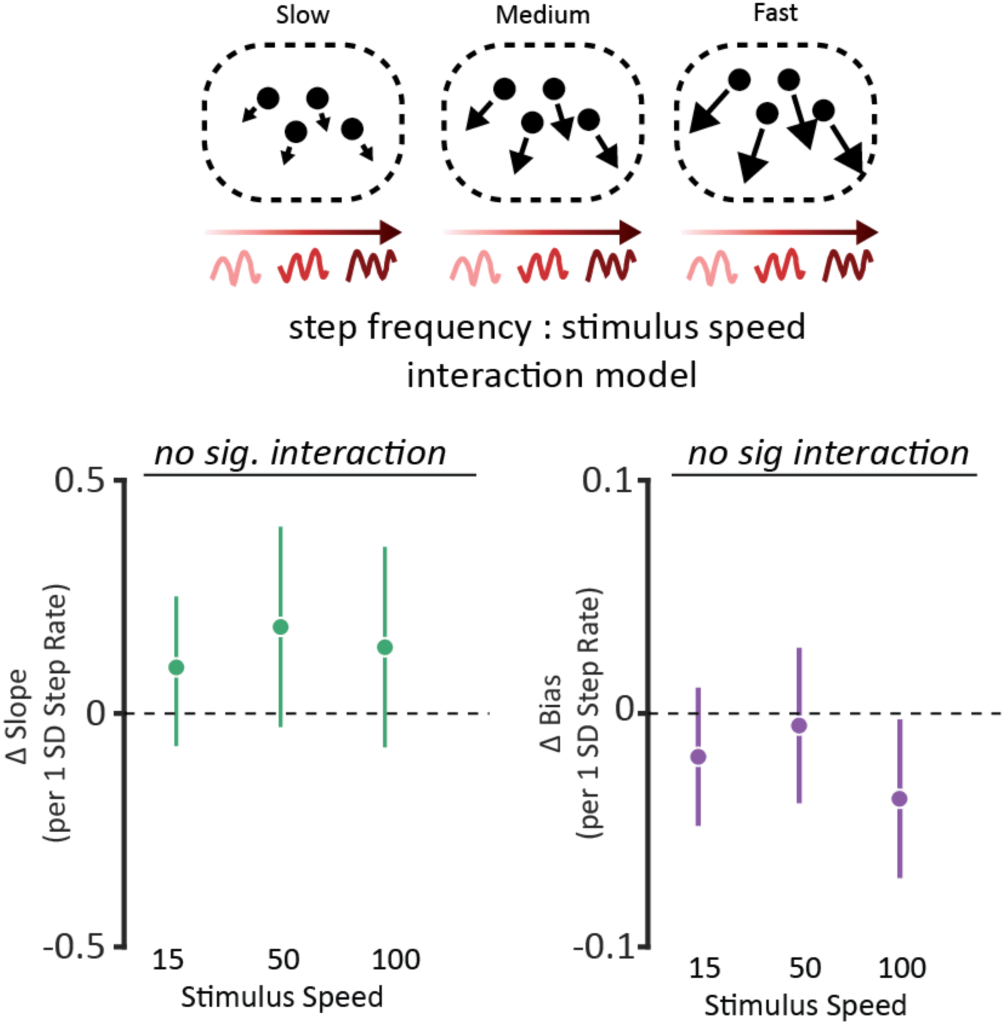
Model weights for the step frequency : stimulus speed interaction model. Model weights for step frequency : stimulus speed interaction model. *Left panel*: effect of step frequency on slope for each stimulus speed. *Right panel*: effect of step frequency on bias for each stimulus speed.

**Supplementary Video 1**

Example set of trials from the experiment, rendered stereoscopically. The stimulus was effectively full-field when viewed through the head-mounted display.

## References

1. Gibson, J.J. (1958). Visually Controlled Locomotion and Visual Orientation in Animals. Ecol. Psychol. 10, 161–176. 10.1111/j.2044-8295.1958.tb00656.x.

2. Lee, D.N. (1980). The optic flow field: the foundation of vision. Philos. Trans. R. Soc. Lond. B. Biol. Sci. 290, 169–179. 10.1098/rstb.1980.0089.

3. Niehorster, D.C. (2021). Optic Flow: A History. -Percept. 12, 20416695211055766. 10.1177/20416695211055766.

4. Parker, A. (2003). In The Blink Of An Eye (Basic Books).

5. Horridge, G.A. (1987). The evolution of visual processing and the construction of seeing systems. Proc. R. Soc. Lond. B Biol. Sci. 230, 279–292. 10.1098/rspb.1987.0020.

6. Egelhaaf, M. (2023). Optic flow based spatial vision in insects. J. Comp. Physiol. A Neuroethol. Sens. Neural. Behav. Physiol. 209, 541–561. 10.1007/s00359-022-01610-w.

7. Nilsson, D.-E. (2022). The Evolution of Visual Roles – Ancient Vision Versus Object Vision. Front. Neuroanat. 16. 10.3389/fnana.2022.789375.

8. Horrocks, E.A.B., Mareschal, I., and Saleem, A.B. (2022). Walking humans and running mice: perception and neural encoding of optic flow during self-motion. Philos. Trans. R. Soc. B Biol. Sci. 378, 20210450. 10.1098/rstb.2021.0450.

9. Gibson, J.J. (1957). Optimal motions and transformations as stimuli for visual perception. Psychol. Rev. 64, 288–295. 10.1037/h0044277.

10. Helmholtz, H. von (1867). Handbuch der physiologischen Optik (L. Voss).

11. von Holst, E., and Mittelstaedt, H. (1950). Das Reafferenzprinzip - Wechselwirkungen zwischen Zentralnervensystem und Peripherie. Naturwissenschaften 37, 464–476. 10.1007/BF00622503.

12. Dauser, H. (1939). Versuche über die Wahrnehmung von Scheinbewegungen durch perspektivische Verschiebungen bei aktiver Kopfbewegung [Experiments on the perception of apparent motion due to perspective shifts during active head movement].

13. Wertheim, A.H. (1994). Motion perception during selfmotion: The direct versus inferential controversy revisited. Behav. Brain Sci. 17, 293–311. 10.1017/S0140525X00034646.

14. Wallach, H. (1987). Perceiving a Stable Environment When One Moves. Annu. Rev. Psychol. 38, 1–29. 10.1146/annurev.ps.38.020187.000245.

15. Wolpert, D.M., and Flanagan, J.R. (2001). Motor prediction. Curr. Biol. 11, R729–R732. 10.1016/S0960-9822(01)00432-8.

16. Durgin, F.H., Gigone, K., and Scott, R. (2005). Perception of visual speed while moving. J. Exp. Psychol. Hum. Percept. Perform. 31, 339–353. 10.1037/0096-1523.31.2.339.

17. Durgin, F.H., and Gigone, K. (2007). Enhanced optic flow speed discrimination while walking: Contextual tuning of visual coding. Perception 36, 1465–1475. 10.1068/p5845.

18. Horrocks, E.A.B., Rodrigues, F.R., and Saleem, A.B. (2024). Flexible neural population dynamics govern the speed and stability of sensory encoding in mouse visual cortex. Nat. Commun. 15, 6415. 10.1038/s41467-024-50563-y.

19. Rowley, D.P., Huk, A.C., and Yates, J.L. (2024). Activity in the peripheral representation within primate V1 is substantially modulated during running. Preprint at bioRxiv, 10.1101/2024.10.10.617723.

20. Cao, L., and Händel, B. (2019). Walking enhances peripheral visual processing in humans. PLoS Biol. 17, e3000511. 10.1371/journal.pbio.3000511.

21. Horrocks, E.A.B., and Saleem, A.B. (2026). Locomotion selectively enhances visual speed encoding in mouse medial higher visual areas. iScience 29. 10.1016/j.isci.2025.114395.

22. Saleem, A.B., Ayaz, A.I., Jeffery, K.J., Harris, K.D., and Carandini, M. (2013). Integration of visual motion and locomotion in mouse visual cortex. Nat. Neurosci. 16, 1864–1869. 10.1038/nn.3567.

23. Blot, A., Roth, M.M., Gasler, I., Javadzadeh, M., Imhof, F., and Hofer, S.B. (2021). Visual intracortical and transthalamic pathways carry distinct information to cortical areas. Neuron 109, 1996–2008.e6. 10.1016/j.neuron.2021.04.017.

24. Duffy, C.J. (1998). MST neurons respond to optic flow and translational movement. J. Neurophysiol. 80, 1816–1827. 10.1152/jn.1998.80.4.1816.

25. Souman, J.L., Freeman, T.C.A., Eikmeier, V., and Ernst, M.O. (2010). Humans do not have direct access to retinal flow during walking. J. Vis. 10, 14. 10.1167/10.11.14.

26. Prsa, M., Gale, S., and Blanke, O. (2012). Self-motion leads to mandatory cue fusion across sensory modalities. J. Neurophysiol. 108, 2282–2291. 10.1152/jn.00439.2012.

27. Pelah, A., Barbur, J., Thurrell, A., and Hock, H.S. (2015). The coupling of vision with locomotion in cortical blindness. Vision Res. 110, 286–294. 10.1016/j.visres.2014.04.015.

28. Thurrell, A. (1998). The influence of non-visual signals of walking on the perceived speed of optic flow.

29. Thurrell, A.E.I., and Pelah, A. (2002). Reduction of perceived visual speed during walking: Effect dependent upon stimulus similarity to the visual consequences of locomotion. J. Vis. 2. 10.1167/2.7.628.

30. Thurrell, A., and Pelah, A. (2005). Matching visual and nonvisual signals: evidence for a mechanism to discount optic flow during locomotion. In Human Vision and Electronic Imaging X, B. E. Rogowitz, T. N. Pappas, and S. J. Daly, eds. (SPIE), p. 434. 10.1117/12.610856.

31. Warren, W.H., and Hannon, D.J. (1988). Direction of self-motion is perceived from optical flow. Nature 336, 162–163. 10.1038/336162a0.

32. Warren, P.A., and Rushton, S.K. (2008). Evidence for flow-parsing in radial flow displays. Vision Res. 48, 655–663. 10.1016/j.visres.2007.10.023.

33. Koenderink, J.J. (1986). Optic flow. Vision Res. 26, 161–179. 10.1016/0042-6989(86)90078-7.

34. Matthis, J.S., Muller, K.S., Bonnen, K.L., and Hayhoe, M.M. (2022). Retinal optic flow during natural locomotion. PLOS Comput. Biol. 18, e1009575. 10.1371/JOURNAL.PCBI.1009575.

35. Rokers, B., Fulvio, J.M., Pillow, J.W., and Cooper, E.A. (2020). Systematic misperceptions of 3-D motion explained by bayesian inference. J. Vis. 18, 23–23. 10.1167/JOV.18.3.23.

36. Czuba, T.B., Rokers, B., Huk, A.C., and Cormack, L.K. (2010). Speed and eccentricity tuning reveal a central role for the velocity-based cue to 3D visual motion. J. Neurophysiol. 104, 2886–2899. 10.1152/jn.00585.2009.

37. Durgin, F.H., Reed, C., and Tigue, C. (2007). Step frequency and perceived self-motion. ACM Trans Appl Percept 4, 5-es. 10.1145/1227134.1227139.

38. Hillman, E.J., Bloomberg, J.J., McDonald, P.V., and Cohen, H.S. (1999). Dynamic visual acuity while walking in normals and labyrinthine-deficient patients. J. Vestib. Res. Equilib. Orientat. 9, 49–57.

39. Grossman, G.E., Leigh, R.J., Bruce, E.N., Huebner, W.P., and Lanska, D.J. (1989). Performance of the human vestibuloocular reflex during locomotion. J. Neurophysiol. 62, 264–272. 10.1152/jn.1989.62.1.264.

40. Al-Yahya, E., Dawes, H., Smith, L., Dennis, A., Howells, K., and Cockburn, J. (2011). Cognitive motor interference while walking: A systematic review and meta-analysis. Neurosci. Biobehav. Rev. 35, 715–728. 10.1016/j.neubiorev.2010.08.008.

41. Woollacott, M., and Shumway-Cook, A. (2002). Attention and the control of posture and gait: a review of an emerging area of research. Gait Posture 16, 1–14. 10.1016/S0966-6362(01)00156-4.

42. Hirasaki, E., Moore, S.T., Raphan, T., and Cohen, B. (1999). Effects of walking velocity on vertical head and body movements during locomotion. Exp. Brain Res. 1999 1272 127, 117–130. 10.1007/S002210050781.

43. Andermann, M.L., Kerlin, A.M., Roumis, D.K., Glickfeld, L.L., and Reid, R.C. (2011). Functional specialization of mouse higher visual cortical areas. Neuron 72, 1025–1039. 10.1016/j.neuron.2011.11.013.

44. Barlow, H.B. (1961). Possible principles underlying the transformation of sensory messages. Sens. Commun., 217–234.

45. Attneave, F. (1954). Some informational aspects of visual perception. Psychol. Rev. 61, 183–193. 10.1037/h0054663.

46. Motyka, P., Akbal, M., and Litwin, P. (2021). Forward optic flow is prioritised in visual awareness independently of walking direction. PLoS ONE 16, e0250905. 10.1371/journal.pone.0250905.

47. Fajen, B.R., and Matthis, J.S. (2013). Visual and Non-Visual Contributions to the Perception of Object Motion during Self-Motion. PLOS ONE 8, e55446. 10.1371/journal.pone.0055446.

48. Barlow, H.B. (1990). A theory about the functional role and synaptic mechanism of visual after-effects. In Vision: coding and efficiency, C. Blakemore, ed. (Cambridge University Press).

49. Barlow, H.B., and Földiák, P. (1989). Adaptation and decorrelation in the cortex. Comput. Neuron, 54–72.

50. Wei, X.-X., and Stocker, A.A. (2015). A Bayesian observer model constrained by efficient coding can explain “anti-Bayesian” percepts. Nat. Neurosci. 18, 1509–1517. 10.1038/nn.4105.

51. Heuer, H. (1987). Apparent motion in depth resulting from changing size and changing vergence. Perception 16, 337–350. 10.1068/p160337.

52. Brenner, E., Van Den Berg, A.V., and Van Damme, W.J. (1996). Perceived motion in depth. Vision Res. 36, 699–706. 10.1016/0042-6989(95)00146-8.

53. Howard, I.P., Fujii, Y., and Allison, R.S. (2014). Interactions between cues to visual motion in depth. J. Vis. 14, 1–16. 10.1167/14.2.14.

54. Rieser, J.J., Pick, H.L., Ashmead, D.H., and Garing, A.E. (1995). Calibration of human locomotion and models of perceptual-motor organization. J. Exp. Psychol. Hum. Percept. Perform. 21, 480–497. 10.1037/0096-1523.21.3.480.

55. Fulvio, J.M., Miao, H., and Rokers, B. (2021). Head jitter enhances three-dimensional motion perception. J. Vis. 21, 1–17. 10.1167/jov.21.3.12.

56. Davidson, M.J., Verstraten, F.A.J., and Alais, D. (2024). Walking modulates visual detection performance according to stride cycle phase. Nat. Commun. 15, 2027. 10.1038/s41467-024-45780-4.

57. Burge, J., and Bonnen, K. (2025). Continuous psychophysics: past, present, future. Trends Cogn. Sci. 29, 481–493. 10.1016/j.tics.2025.01.005.

58. Schütt, H.H., Harmeling, S., Macke, J.H., and Wichmann, F.A. (2016). Painfree and accurate Bayesian estimation of psychometric functions for (potentially) overdispersed data. Vision Res. 122, 105–123. 10.1016/j.visres.2016.02.002.

